# Signal and Measurement Considerations for Human Translation of Diffuse in vivo Flow Cytometry

**DOI:** 10.1101/2022.03.23.485369

**Authors:** Fernando Ivich, Joshua Pace, Amber L. Williams, Malcolm Shumel, Qianqian Fang, Mark Niedre

## Abstract

**Significance:** Diffuse in vivo flow cytometry (DiFC) is an emerging technology for fluorescence detection of rare circulating cells directly in large deep-seated blood vessels in mice. Because DiFC uses highly scattered light, in principle it could be translated to human use. However, an open question is whether fluorescent signals from single cells would be detectable in human-scale anatomies.

**Aim:** Suitable blood vessels in a human wrist or forearm are at a depth of approximately 2-4 mm. The aim of this work was to study the impact of DiFC instrument geometry and wavelength on the detected DiFC signal and on the maximum depth of detection of a moving cell.

**Approach:** We used Monte Carlo simulations to compute Jacobian (sensitivity) matrices for a range of source-detector separations and tissue optical properties over the visible and near infrared (NIR) spectrum. We performed experimental measurements with three available versions of DiFC (488 nm, 640 nm, and 780 nm), fluorescent microspheres, and tissue mimicking optical flow phantoms. We used both computational and experimental data to estimate the maximum depth of detection at each combination of settings.

**Results and Conclusions:** For the DiFC detection problem, our analysis showed that for deep-seated blood vessels, the maximum sensitivity was obtained with NIR light (780 nm) and 3 mm source-and-detector separation. These results suggest that - in combination with a suitable molecularly targeted fluorescent probes - circulating cells and nanosensors could in principle be detectable in circulation in humans.

## 1. Introduction

Diffuse in vivo flow cytometry (DiFC)^1,2^ is an emerging technique that uses diffusely scattered light to continuously and non-invasively detect and count fluorescence-labeled (and fluorescent protein expressing) cells in the blood of small animals. DiFC uses optical fiber bundles (“probes”) to generate and measure laser induced fluorescent light from individual cells moving in blood vessels, for example in the mouse tail artery (***Fig. 1a***)^3^. A unique property of DiFC is that it allows sampling of the full peripheral blood volume of a mouse in about 15 minutes. A major application is therefore non-invasive enumeration of rare circulating tumor cells (CTCs), which have been found to be instrumental in hematogenous metastasis but typically number fewer than 100 cells per mL of peripheral blood. We previously used DiFC to detect rare CTCs in xenograft models^4,5^ and observe changes in CTC numbers over time^6^. We also used DiFC with engineered optical sensors that circulate in the blood to measure systemic sodium levels^7^.

**Fig. 1.**
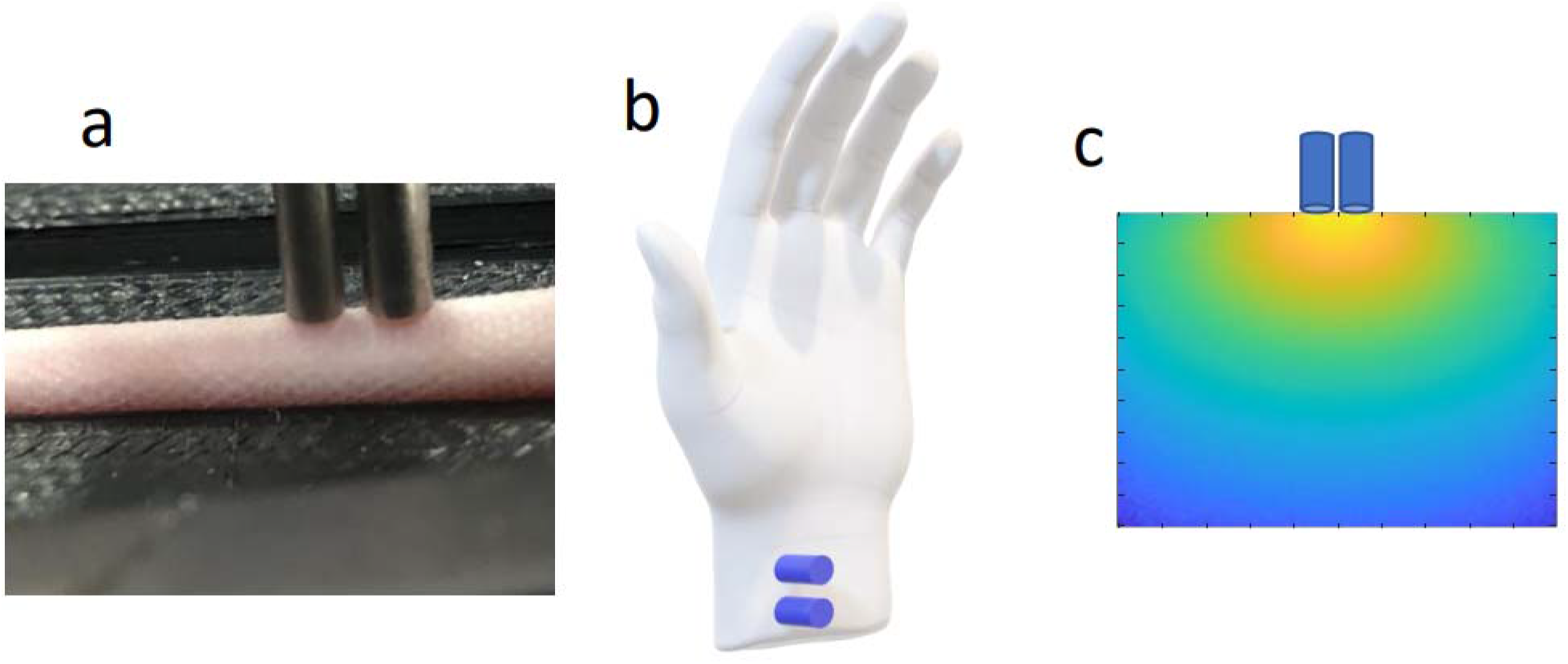
(a) Photograph of DiFC optical probes on the ventral surface of a mouse tail. We use DiFC to detect rare fluorescently-labeled CTCs in the ventral caudal artery of mice. (b) Conceptual diagram of DiFC use in human wrists, where blood vessels are 2-4 mm in depth. (c) Qualitative example sensitivity profile (Jacobian matrix) for a 3-mm source-detector separation.

Because DiFC uses diffuse light and works in an epi-illumination (as opposed to trans-illumination) geometry, in principle it could be used in larger species including humans. If feasible, DiFC could provide a new method to continuously enumerate rare CTCs ^8–10^ directly *in vivo* without having to draw and analyze blood samples^11^. Fluorescent labeling of CTCs for DiFC could be achieved using cancer-specific fluorescence contrast agents such as those in development for fluorescence guided surgery^12,13^. Our lab showed that it is feasible to label CTCs directly in mice with a small molecule folate targeted fluorescent probe (EC-17), and detect them externally with our DiFC system^14^.

However, translation of DiFC to humans would also entail detecting light from deeper-seated blood vessels compared to those in mice. Specifically, we have used DiFC on arteries in the tail or leg in mice which are approximately 0.75-1 mm in depth. Suitable candidate blood vessels in a human include the radial artery or vein in the human wrist which are about 2-4 mm in depth^15^ (***Fig. 1b***). An open question is therefore whether single cells would be detectable with DiFC since the fluorescent signal is expected to be significantly attenuated due to light scatter and absorption (***Fig. 1c***). In addition, the ratio of the signal from a single cell to the background (non-specific) tissue autofluorescence is expected to be lower at deeper blood vessel depths.

As described in more detail below, our small animal DiFC design uses an integrated multi-fiber probe, where the source and detector fiber positions are physically adjacent (0.3 mm center-to-center separation). Previous work in the near-infrared spectroscopy (functional NIRS) fields have extensively studied the effect of different fiber probe geometries on the optical sensitivity of human tissue^16^. In particular, NIRS and diffuse optical tomography (DOT) frequently use larger source-detector separations to probe deeper tissue volumes^17–19^, for example in the brain^20,21^. However, it is unclear to what degree this approach is applicable for the specific DiFC detection problem. It is also broadly understood that red and NIR light undergoes less attenuation in tissue than visible light^22,23^.

The purpose of this work was to use Monte Carlo (MC) photon transport simulations and experimental optical phantom models to i) study the effect of laser (and fluorophore) wavelengths, and light source and detector separation on the detection problem in DiFC, and ii) to assess whether in principle CTCs could be detectable in appropriate blood vessels in humans.

## 2. Materials and Methods

### 2.1 Diffuse in vivo Flow Cytometer (DiFC)

DiFC instrumention has been described in detail by our team previously^1,4,6,7^. Thus far we have developed blue-green^4^ (488 nm laser), red ^1^ (640 nm laser), and NIR (780 nm laser) versions, designed to work with different widely-used fluorophores and fluorescent proteins. Briefly, DiFC uses laser light which is coupled into an optical fiber to illuminate a sample surface (i.e., skin). As fluorescently-labeled cells pass through the field-of-view, they emit fluorescent light which is detected with a separate set of 8 detection fibers. Each fiber tip has miniaturized integrated filters to minimize leakage of laser light into the collection fiber. The output is filtered and detected with photomultiplier tubes (PMTs).

In normal DiFC operation^1,4^, the source and detector fibers are assembled in a bundle, with center-to-center separation of 0.3 mm (***Fig. 2a)**.* However, we are interested in the impact of wider source-and-detector fiber separation on cell detection depth and sensitivity^15^. In this case, we used two separate fiber bundles – one as the excitation source and the second is for collection of fluorescent light only. These are separated by a source-and-detector distance of 3 mm or greater.

**Fig. 2.**
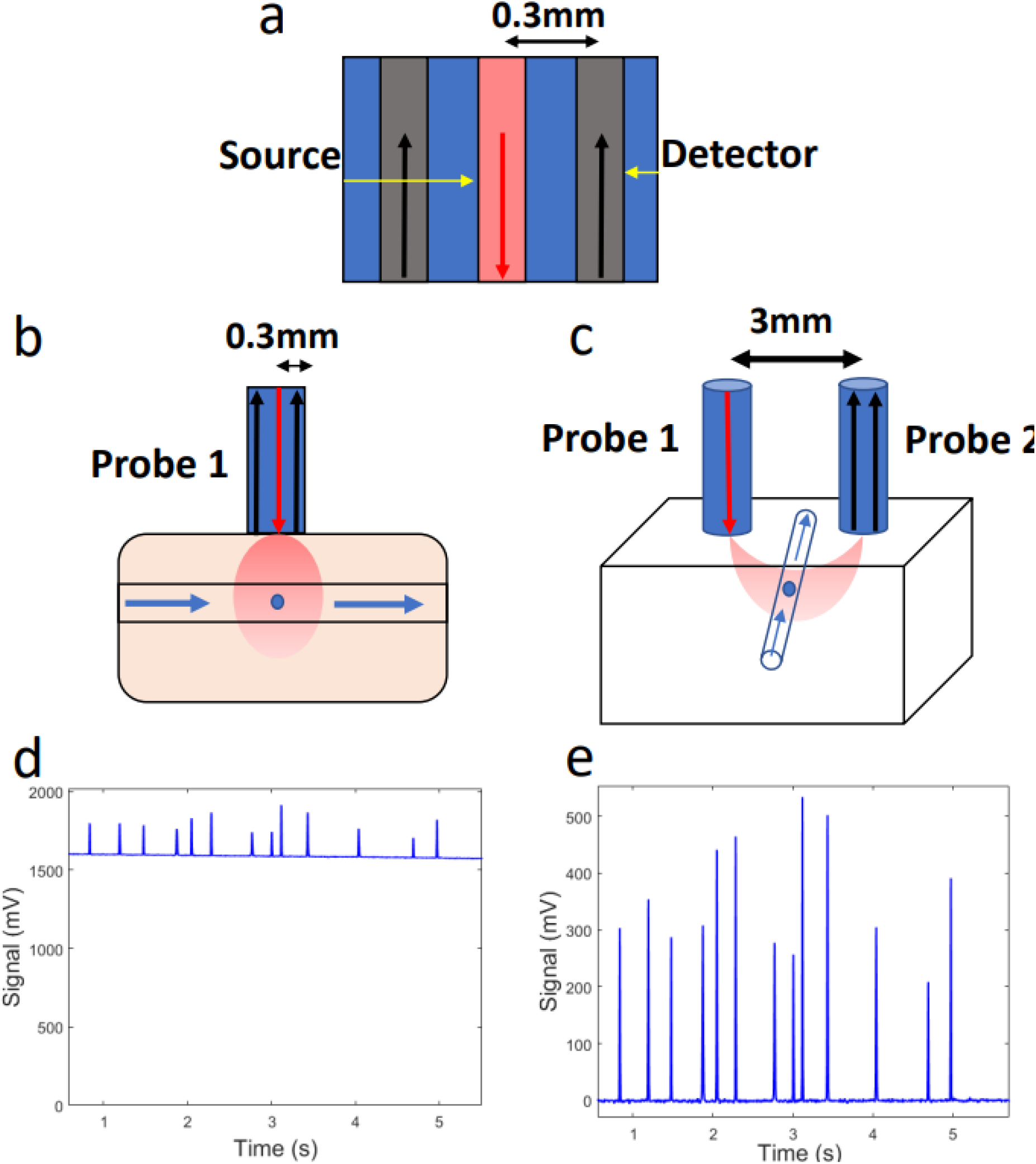
Diffuse in vivo Flow Cytometer (DiFC). (a) Diagram of DiFC fiber probe showing the detector fibers in black and the source fiber in red. (b) 0.3 mm source-detector separation configuration for DiFC. The DiFC fiber probe is showing the laser source (red arrow) and detector (black arrows) separated by 0.3 mm. (c) Two DiFC fiber probes separated by 3 mm. Fiber probe 1 has a light source (red arrow) and fiber probe 2 has photodetectors (black arrows) separated by 3 mm. (d) Non-background-subtracted sample data showing cell detections as peaks. (e) Background-subtracted sample data showing the same cell detections.

### 2.2 Monte Carlo Light Transport Simulations

Monte Carlo (MC) simulations of photon propagation in 3D tissue volumes were performed using Monte Carlo eXtreme (MCX) software^24,25^. MCX is an open-source MC simulator that accelerates computational efficiency via parallel processing using a graphic processing unit (GPU). Simulations involved using a cubic homogeneous 5 ×5 ×5 cm^3^ volume representing a portion of a human arm, with an isotropic voxel size of 250 μm. The optical properties for a selective set of wavelengths within the visible and near infrared (NIR) spectrum were chosen from the literature^26^. The refractive index and anisotropy coefficient were chosen to be 1.37 and 0.9, respectively, for all tested wavelengths. **Table 1** shows the optical properties used for the studies here. Although there is of course a wavelength red-shift between excitation and emission laser light, this effect is negligible compared to the uncertainties in the values (see discussion below)^26,27^. For each simulation, a total of 10^8^ photon packets were simulated with a time gate of 5 ns.

**Table 1.**
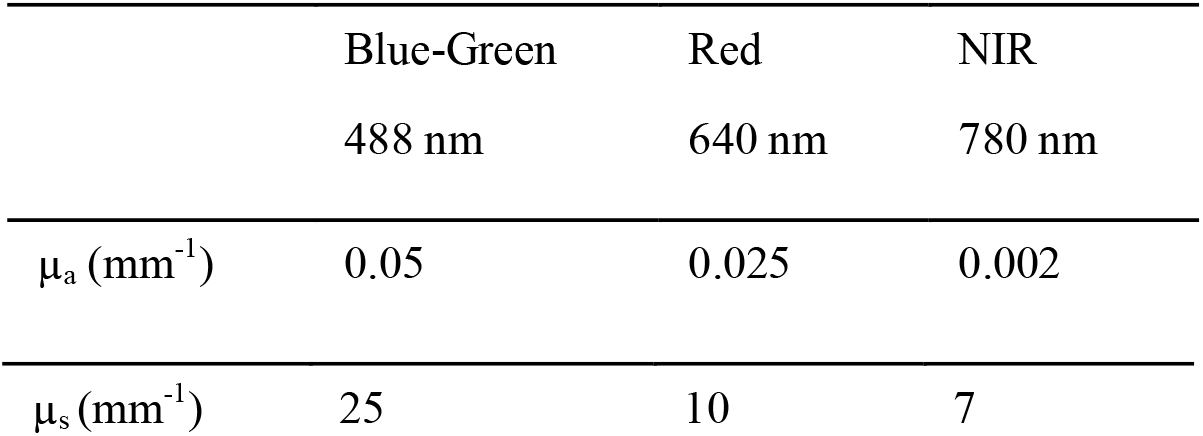
Absorption (μ_a_) and scattering (μ_s_) coefficients used for the DiFC Monte Carlo simulations. The anisotropy coefficient, g, and the index of refraction, n, were 0.9 and 1.37, respectively, in all cases^25^.

|  | Blue-Green | Red | NIR |
| --- | --- | --- | --- |
|  | 488 nm | 640 nm | 780 nm |
| $\mu_a \text{ (mm}^{-1}\text{)}$ | 0.05 | 0.025 | 0.002 |
| $\mu_s$ (mm<sup>-1</sup>) | 25 | 10 | 7 |

Each element of the absorption sensitivity matrix *W(r; r_s_, r_d_*) reflects the magnitude of the detection signal change for a given source (located at *r_s_*) detector (located at *r_d_*) pair for a unitary μ_a_ perturbation at a given location *r*^28^. This was computed via the adjoint Monte Carlo (aMC) method^28^, by multiplying the forward solution of the radiative transfer equation of the light source (source-fluence) by the forward solution of the radiative transfer equation at the detector position (detector-fluence). Simulations were performed for a range of source-and-detector separations (SDS) ranging from 0.3 mm to 12 mm on the surface of the volume. The extracted voxel-based sensitivity values from the simulations were taken from the middle point between source and detector along the depth of the simulated tissue, as shown in ***Figs. 2b,c***.

### 2.3 Contrast (Signal to Background Ratio) Estimation

To explore theoretical limits of detection, we estimated the sensitivity ratio between a fluorescent peak and the background autofluorescence as follows. Following the first Born approximation, this linearized formulation follows the equation U_fl_(r_s_,r_d_) = W(r; r_s_, r_d_)× η(r). Where *U_fl_* is the measured signal between a source (*r_s_*) and detector (*r_d_*), *W* is the sensitivity matrix as described above, and *η(r*) is the fluorophore concentration as a function of position in the media.

To estimate the signal for a cell detection (peak) *U_fl-pk_*(*r_s_,r_d_*) at depth *d*, we assumed η(r) = ηcell for r = (0,0, d), and η(r) = 0 elsewhere. Note that the diameter of a cell is much smaller than the voxel size used here. To estimate *η_cell_*, we used experimental DiFC measurements as detailed below.

To estimate the background autofluorescence signal *U_af_*(*r_s_,r_d_*), we made the simplifying assumption that autofluorescence was constant throughout the media *η_af_*. This was also estimated using experimental measurements. Note that both *η_cell_* and *η_af_* were assumed to be different for the 488, 640, and 780 nm systems.

### 2.4 Optical Phantom Models in vitro

To experimentally test the relationships between target depth, source-detector separation distance, and wavelength, we used a tissue mimicking optical phantom made of high-density polyethylene. We have previously shown that this material has optical properties similar to biological tissues^1^. The phantom has drilled holes at different depths ranging from 0.75 mm to 4 mm deep where we thread microbore Tygon tubing (TGY-010-C, Small Parts, Inc., Seattle, Washington) to simulate a blood vessel. The tubing is connected to a syringe pump (70-2209, Harvard Apparatus, Holliston, Massachusetts) where we flow fluorescent microspheres to mimic fluorescently labeled cells^1,4^, or optical sensors. We suspended the fluorescent microsphere solution in phosphate buffer saline (PBS) at a concentration of 10^3^ microspheres per milliliter and flowed them at a rate of 50 μl per minute. We used Flash Red reference intensity 5 (FR5; Bangs Laboratories Inc., Fisher, Indiana), Dragon Green reference intensity 5 (DG5; Bangs), and Jade Green high intensity (Spherotech Inc., Lake Forest, Illinois) microspheres for red, blue-green and NIR wavelengths, respectively. We showed previously that these approximate the fluorophore brightness of a well-labeled cell^1,14^.

**In *Fig. 2***, we include a schematic of the DiFC fiber probe showing the source fiber and the detector fibers (***Fig. 2a***). Example configurations for 0.3 mm and 3 mm SDS are shown in ***Figs. 2b*** and ***2c*** respectively. The tubing is placed in the middle of the two probes, running in a perpendicular direction. ***Fig. 2c*** shows example measured data, and ***Fig. 2d*** shows the data after mean background subtraction. When increasing the source-and-detector separation (SDS) separation, we also increased the PMT sensitivity gain to approximately match the background autofluorescence. The laser power at the sample surface was 20 mW for all experiments.

### 2.5 DiFC Data Analysis

DiFC data was analyzed as described previously^4^. We repeated each run 5 times for every wavelength combination, SDS, and microsphere-depth. Peaks that exceeded a threshold of five times the noise (standard deviation) were counted as a detection event, and the amplitude was recorded. We also used control runs (N=5) with PBS only (no microspheres) to determine the instrument false alarm rate (FAR).

## 3. Results

### 3.1 Monte Carlo Simulations

Sensitivity matrices were computed for 488, 640 and 780 nm excitation wavelengths and for different SDS values as summarized in ***Fig. 3***. ***Figs. 3a,c,e*** show example sensitivity matrices for SDS of 0.3 mm, 3 mm, and 6 mm, respectively. ***Figs. 3b,d,f*** show the sensitivity as a function of depth (up to 5 mm) at the center line between the fibers at 488 nm, 640 nm, and 780 nm for the same SDS. These data are normalized to the maximum sensitivity value along the midline between source and detector, for all combinations, in this case 0.3 mm SDS and 488 nm wavelength. The effects of SDS on the depth sensitivity of tissue are summarized in ***Figs. 3g,h***. As expected, these results showed that the depth of maximum sensitivity increased with increased SDS (***Fig. 3g***). While these data nominally suggest that deeper cell sensitivity could be achieved using larger SDS, we note that the absolute value of the sensitivity also decreased strongly (by ~3 orders of magnitude) for larger SDS (***Fig. 3h)***. Likewise, the absolute value of sensitivity generally decreased with shorter wavelengths due to higher light attenuation (***Fig. 3h***). The implications of these for the DiFC detection problem are discussed in more detail in section 3.3 below.

**Fig. 3.**
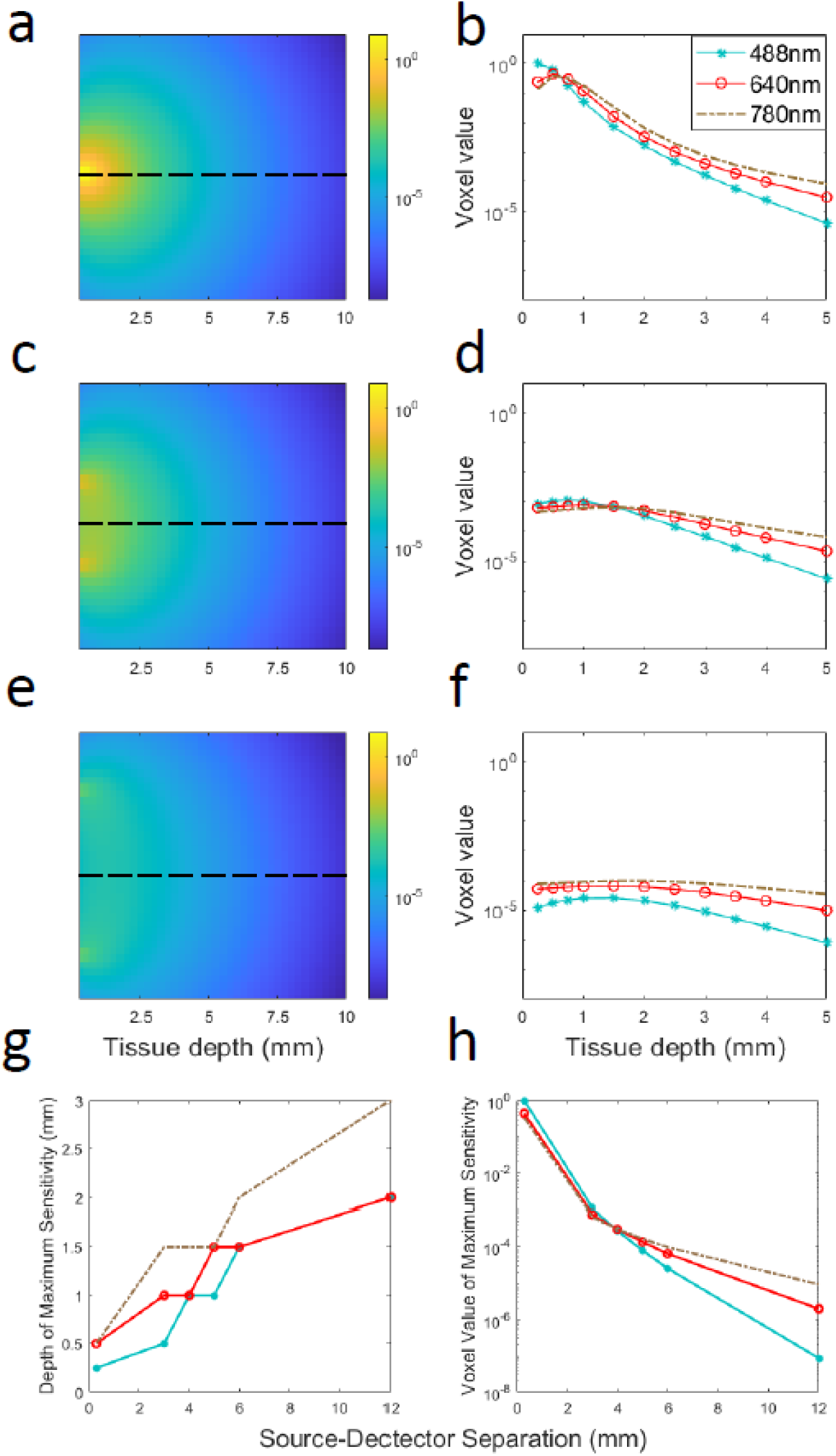
Monte Carlo simulations of photon propagation in a tissue-mimicking domain with optical properties as in ***Table 1***. (a) Example sensitivity matrix for 0.3 mm SDS and 488 nm wavelength. (b) sensitivity depth profile for SDS of 0.3 mm for 488, 640 nm, and 780 nm. (c,d) example sensitivity matrix and depth profiles for a 3 mm SDS, and (e,f) for a 6 mm SDS. (g) Depth of maximum sensitivity for different SDS. (h) Normalized (to the maximum along the midline) maximum sensitivity value for different SDS.

### 3.2 Measurements in Phantoms In Vitro

We next used our three DiFC systems to experimentally test the same relationships in a tissue-mimicking flow phantom model^1,4^. ***Fig. 4*** shows the normalized MC sensitivity calculations and experimental measurement of mean fluorescent microsphere peak amplitudes for 0.3 mm and 3 mm SDS. Specifically, ***Fig. 4a*** shows the mean peak amplitude of DG5 microspheres (symbols) measured with DiFC with MC sensitivity calculations (solid line) for 488 nm and 0.3 mm SDS. ***Fig. 4b***, shows the same with 3 mm SDS. Likewise, ***Figs. 4c,d*** show the same relationships for FR5 microspheres and 640 nm MC calculations, and ***Figs. 4e,f*** show the same relationships for Jade Green microspheres and 780 nm MC calculations. Here, each solid data point shows the average, and standard deviation from N = 5 measurements. For all wavelengths, the 0.3 mm SDS yielded greater signal amplitude at shallow target depths. We observed a small increase in amplitude for 780 nm for deeper targets for 3 mm SDS compared to 0.3 mm. The maximum detection depth was 2 mm for 488 nm and 640 nm, and 3.5 mm for 780 nm. For peaks of sufficiently low amplitude (near the instrument noise floor around 5 mV), it is likely that some microspheres were simply below the detection threshold of the system. To better illustrate this, we plotted the normalized (to maximum) detection count rate (per minute) for 0.3 mm and 3 mm SDS, in ***Figs. 5a,b***, respectively. As shown, the normalized count rate drops significantly at larger depths, and is more pronounced for lower wavelengths.

**Fig. 4.**
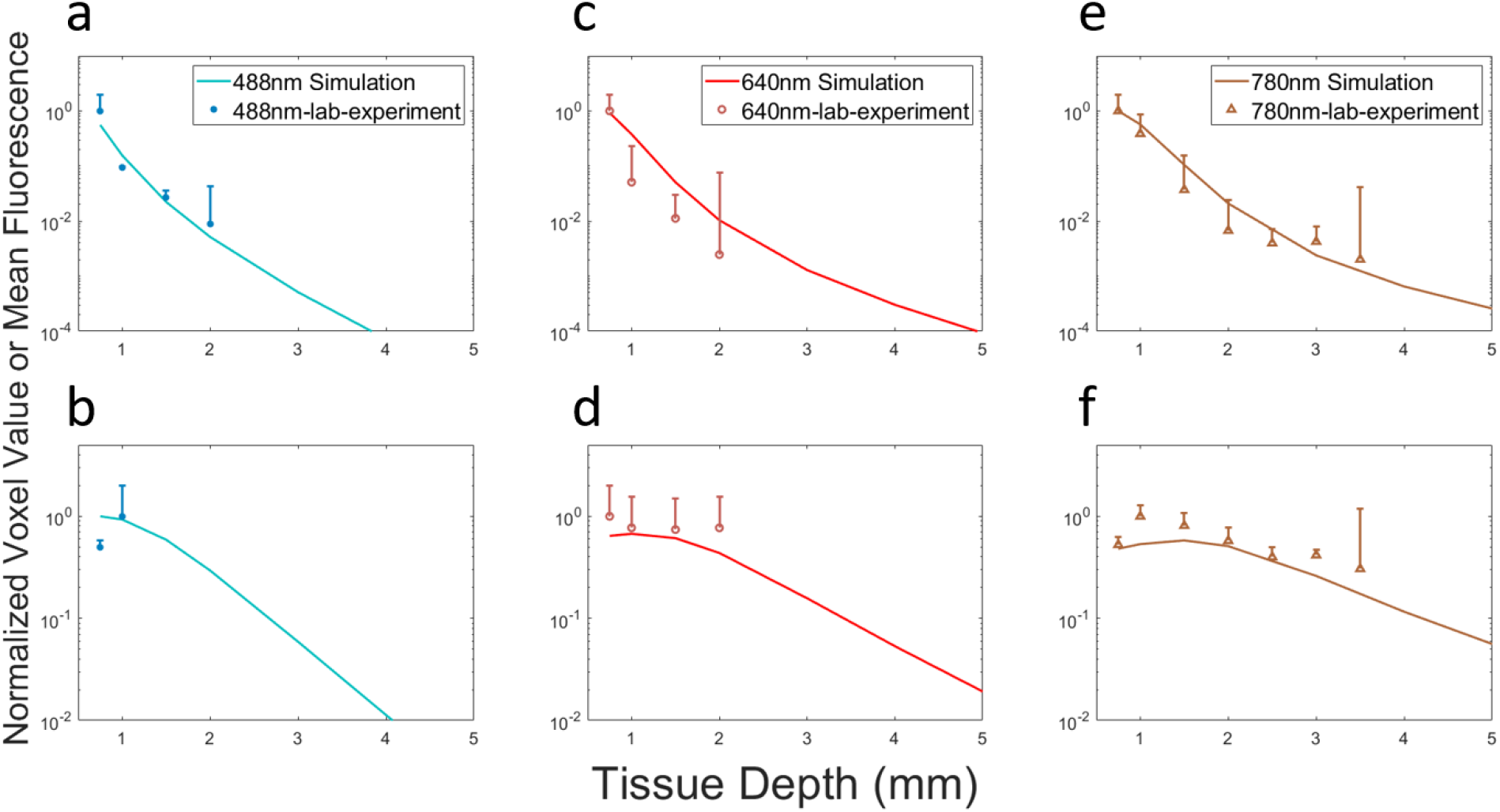
Optical phantom measurements for 0.3 mm (a,c,e) and 3mm (b,d,f) separations for 488, 640 and 780 nm wavelengths, respectively. The normalized MC simulated sensitivity and experimental measurements of mean DiFC peak intensity of microspheres for each are shown. The error bars are the standard deviation of N=5 repeats in each case.

**Fig. 5.**
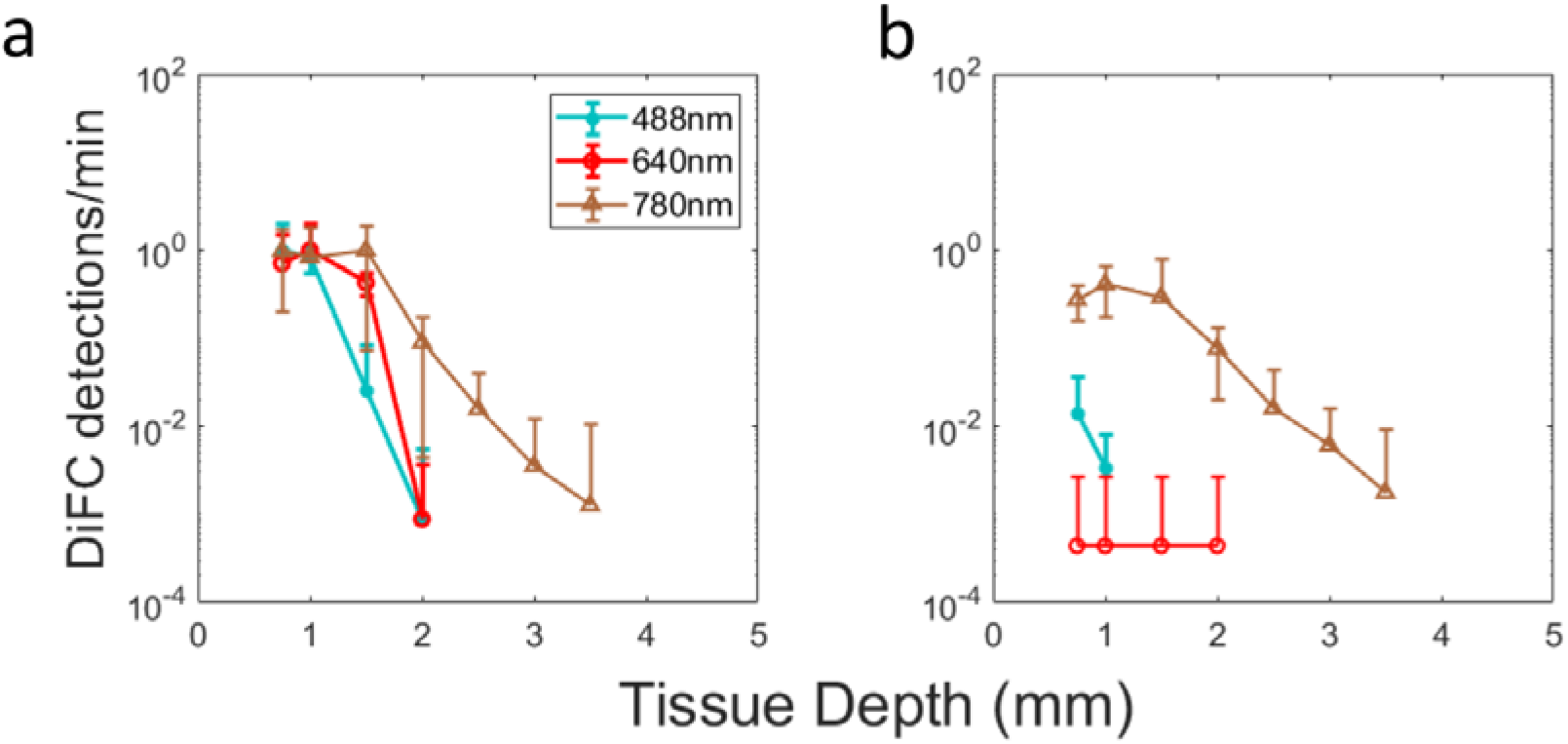
Normalized microsphere detections per minute for nominally identical suspension concentrations for (a) 0.3 mm and (b) 3 mm SDS.

### 3.3 Contrast to Background Analysis

The DiFC detection problem relies on measurement of a small fluorescence signal from a single moving cell, on top of a larger non-specific background autofluorescence signal. This background is approximately constant over the timescale of a detected CTC peak (seconds). The magnitude of this autofluorescence varies with the laser wavelength and the type of tissue (e.g., mouse strain) in the experiment. While we subtract the *mean* of this background in data processing (as in ***Figs. 2c,d*)**, the additive *noise* cannot be subtracted. The ratio of the expected peak amplitude from a single cell to this background noise therefore defines the lower level of detection sensitivity.

We can estimate this threshold for arbitrary SDS for each wavelength using our experimentally measured (phantom) data above and MC sensitivity functions. We estimated an average background autofluorescence concentration ***η_af_*** in each case, which we assumed was homogeneous throughout the media to give the mean autofluorescence signal. We modeled additive Gaussian noise, which based on our measurements with our existing DiFC prototypes, was a percentage of the average PMT signal output. Assuming a baseline (autofluorescence) signal output of 10% the maximum, the noise was equal to 0.2% of the background amplitude. We also estimated the average fluorescence concentration of a cell ***η_cell_*** (which we assume was smaller than a voxel) to calculate the peak amplitude. We assumed that the minimum detectable peak was 5 times the noise (signal to noise ratio = 13.9 dB). This threshold yielded a FAR of 0.01 false positive detections per minute over all conditions tested. The results are summarized in ***Fig. 6***.

**Fig. 6.**
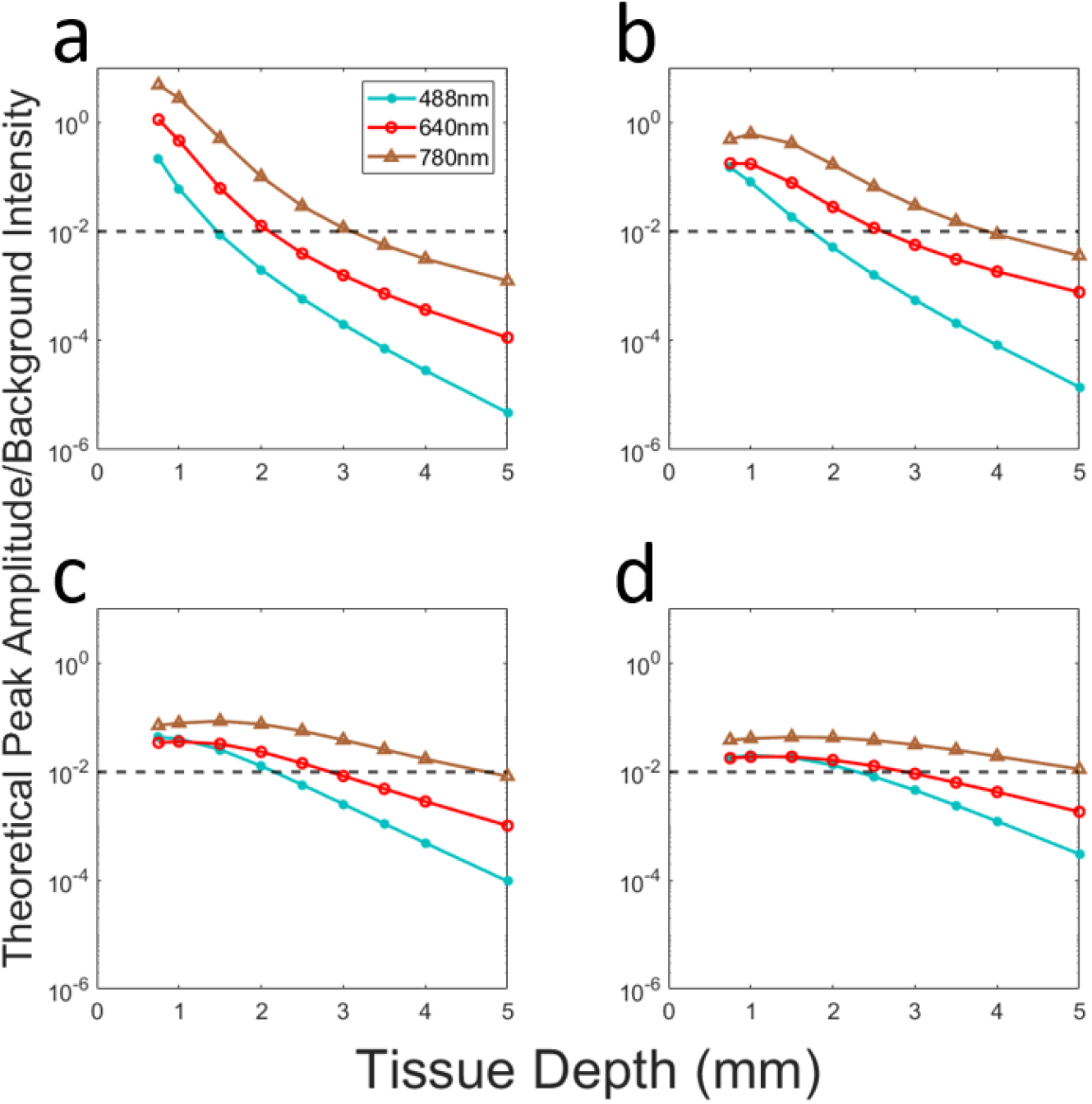
Estimated signal to background ratios for red, blue-green, and NIR light. Horizontal lines represent the detection limit of 13.9 dB for (a) 0.3 mm, (b) 1 mm, (c) 3 mm, and (d) 5 mm SDS, respectively.

As shown, the results are generally consistent with our experimental measurements. First, as expected, 780 nm yielded highest peak to background ratio in all cases, followed by 640 nm, and 488 nm. Second, use of larger SDS in general allowed detection of cells at larger depths particularly at longer wavelengths. Assuming cells were approximately 1 mm in depth (as is the case with our pre-clinical mouse experiments^4^), cells were expected to be detectable with all three wavelengths. If cells are expected to be 2 to 4 mm in depth (as expected in in humans) then larger SDS are expected to yield higher peak-to-background ratios for 780 nm DiFC. Specifically, for 2-, 3-, and 4-mm deep cells, SDS of 1, 2, and 5 mm are expected to yield maximum sensitivity, and 2 mm source-and-detector separation are expected to have best median sensitivity over the 2-4 mm depth range.

## 4. Discussion and Conclusions

We previously reported the use DiFC exclusively in mice^1,2,4,29^, although an open question is whether DiFC could work in humans. The purpose of this work was to study whether tissue optics would permit detection of single cells in blood vessels 2-4 mm deep in diffuse tissue such as the radial^15^ or ulnar artery or superficial facial arteries^30^. Moreover, the goal was to determine which SDS configuration and wavelengths of light would be most appropriate. Although the effect of fiber probe design has been widely studied for diffuse optical tomography applications^17–19^, we are unaware of any other theoretical and experimental study to explore the specific problem of diffuse fluorescence measurement from a single cell in bulk tissue.

Beyond DiFC, there are other technologies in development for optical detection of CTCs in vivo^5^, including photoacoustic methods^31, 32^ or intravital microscopy^33–35^ with similar considerations for light attenuation in biological tissue. As noted, because DiFC works with diffuse light, it is expected to allow significantly greater depth-of-penetration compared to intravital confocal microscopy-based methods and is uniquely suited for human translation.

The MC studies here made the simplifying assumptions that the tissue was homogenous, and that the optical properties were the same for excitation and emission light. It also assumes that the background non-specific autofluorescence (particularly at superficial tissue depths) is approximately constant over the timescale of a CTC detection (seconds), which is valid in our prior experimental measurements. As shown, despite these assumptions our theoretical and experimental results were in good general agreement, and also consistent with our prior experimental work with DiFC in mice. This said, we tested how sensitive our conclusions were to the assumed optical properties by varying μ_a_ and μ_s_ by up to 50% for each combination. The estimated peak amplitude to background ratios (analogous to ***Fig. 6***) for varying optical properties for 0.3 and 3 mm separation are shown in ***Figs. 7a,b***. As shown, varying the optical properties resulted in a change to the maximum estimated detection depth of 0.5 mm or less.

**Fig. 7.**
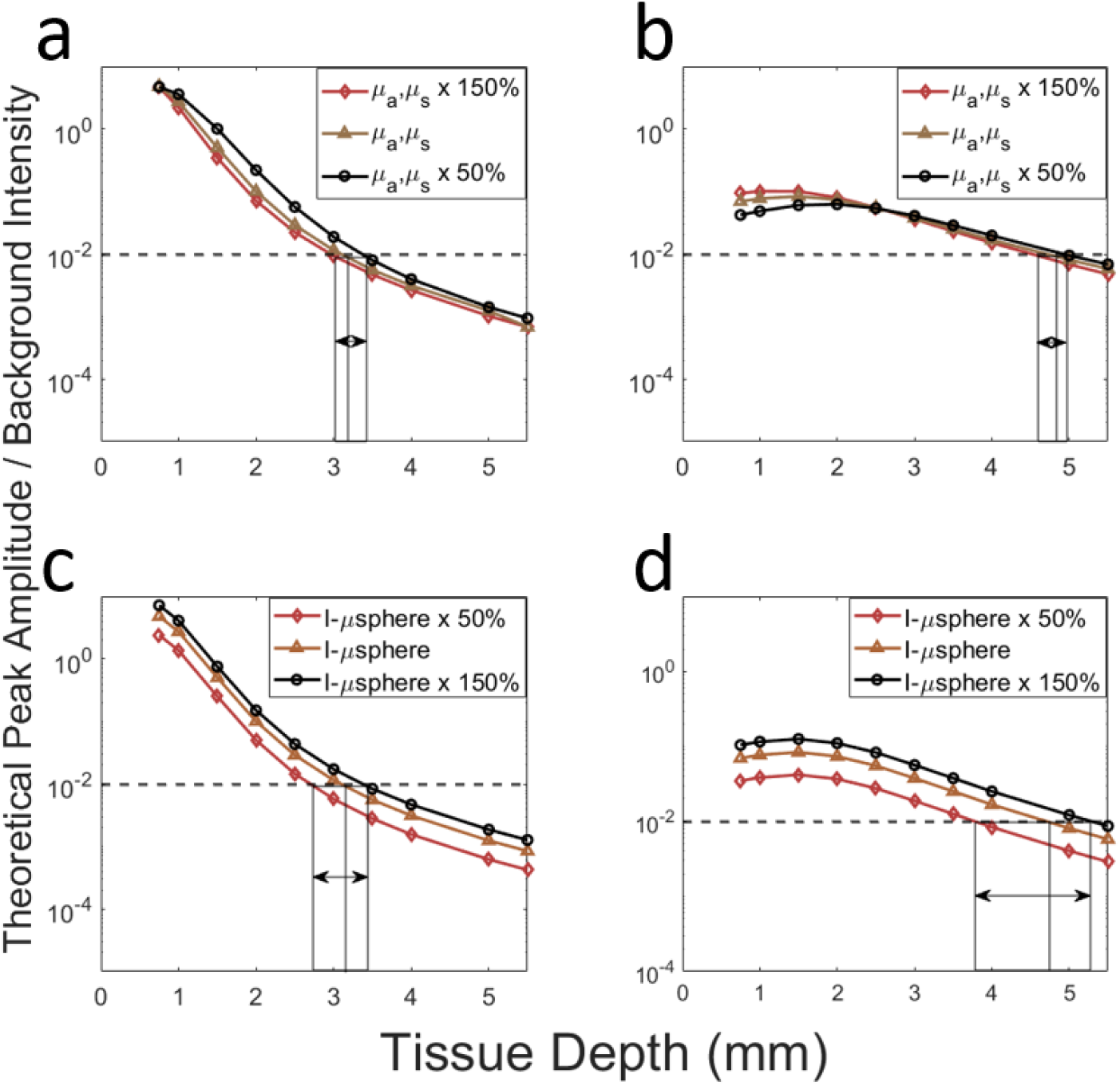
(a,b) Estimated peak to background ratios for 780 nm light assuming 50% higher and 50% lower in optical properties for (a) 0.3 mm SDS and (b) 3 SDS, respectively. Estimated peak to background ratios assuming cell brightness 50% brighter 50% dimmer for (c) 0.3 mm, and (d) 3 mm SDS, respectively.

Likewise, we made the simplifying assumption that fluorescent microspheres used here approximated the brightness of a well-labeled (or fluorescent protein expressing) cell. This is generally consistent with our previous work^1^. In practice this brightness is a complex combination of fluorophore uptake by each cell^36^, extinction coefficient, quantum yield, and emission spectra match with the instrument optical filters, all of which would also naturally affect the maximum depth of detection in DiFC. In order to test this, we calculated the effect of detection depth if cells were either 50% brighter or 50% dimmer (***η_cell_)*** as summarized in ***Fig. 7c,d***. Increasing or decreasing the brightness may affect the maximum detection depth by as much as 1 mm. Likewise, our assumption of detectability relies on an assumption of instrument noise (0.2% of PMT output). In practice, noise may be higher in living organisms due to, for example, motion or photoplethysmography (PPG) artifacts^37^, which in our experience is more pronounced in our green fluorescent protein (GFP) compatible DiFC system. Likewise, the estimated detection limit could be improved with alternate signal processing approaches, reduction in instrument noise, or use of a lower threshold.

Overall, these studies showed two main practical insights for DiFC. First, for cases where cells are presumed to be at a shallow (<2 mm) depth, the use of a small (0.3 mm) SDS resulted in the highest sensitivity and allow use of blue-green, red, and NIR wavelengths. Unsurprisingly, blue-green optical properties performed the worst, both due to the high non-specific tissue autofluorescence and the attenuation of light in the tissue. However, in small animal geometries (~1 mm deep blood vessels), the signal to noise ratio and experimental analysis (along with our prior experience) demonstrate that cells should be detectable. Because of the broad availability of GFP expressing cells, 488 nm DiFC is valuable in pre-clinical research. However, these data indicate that it is unlikely to be practical in translation to humans unless extremely bright fluorescent labeling can be achieved.

Second, for cases where cells are presumed to be present at larger depths (2-4 mm, as we expect in humans), SDS between 1-5 mm performed the best, with good general median coverage of the depth range observed for 3 mm separation. As expected, the highest detected signal to noise ratio for a given depth was obtained using NIR wavelengths. Most of the recent class of fluorescence guided surgery molecular contrast agents for cancer are NIR^13,38^, some of which may ultimately be used as a contrast agent for DiFC. Likewise, circulating fluorescent sensors^7,39^ can be engineered to emit light in NIR wavelengths.

In summary, the most promising DiFC instrument geometry for potential human translation is with a source-and-detector separation around 3 mm and using NIR fluorescent contrast agents. Experimental and computational analyses presented here suggest that this configuration should allow interrogation of human blood vessels located 2 to 4 mm under the skin, which is suitable for several major blood vessels that carry large circulating blood volumes. Evaluation and characterization of NIR contrast agents for CTCs is an ongoing area of work in our group^14^.

## Acknowledgements

This work was funded by the National Institutes of Health (R21CA246413, R01EB024186, R01-GM114365 and R01-GM114365).

## Code, Data and Materials Availability

Matlab code and data for the Monte Carlo simulations, the laboratory experiments and the generation of the figures can be found in Github open-source library (https://github.com/NiedreLab/Ivich_depth_sensitivity_2022).

**Fernando Ivich** received his BS and MS degrees in biomedical engineering from the University of Arizona in 2017 and 2019, respectively. He is a PhD candidate in bioengineering from Northeastern University.

**Mark Niedre** received his PhD from the University of Toronto in the department of medical physics in 2004. He is a professor of bioengineering at Northeastern University and a senior member of SPIE.

